# The influence of similarity, sensitivity and bias on letter identification

**DOI:** 10.1101/2025.04.20.649714

**Authors:** Hatem Barhoom, Mahesh R. Joshi, Gunnar Schmidtmann

**Affiliations:** Eye & Vision Research Group, School of Health Professions, University of Plymouth, UK; Islamic University of Gaza, P.O. Box 108, Gaza, Palestine; Centre for Visual Science, University of Rochester, NY, USA

**Keywords:** bias, similarity, sensitivity, noisy template model

## Abstract

Previous studies have demonstrated that bias, sensitivity and similarity between letters are causes of errors in letter identification. However, these factors and their relative contribution in letter identification have not been investigated extensively. Our previous model (noisy template model) was devised to calculate the effect of bias and sensitivity in letter identification task. In the current study, we used the method of constant stimuli to measure letter acuity for Sloan letters at an eccentricity of 7 deg from fixation (temporal visual field). Similar to our previous work, we devised an tested a variety of models to estimate the joint role of bias and sensitivity, but extended our model to also incorporate the similarity between letters.

Modelling results showed that bias is the major factor in determining the pattern of total, correct and incorrect responses in letter identification. Furthermore, the joint effect of similarity and bias was found to be higher than the joint effect of either bias and sensitivity or similarity and sensitivity in shaping the pattern of overall responses in letter identification. Incorporating the similarity factor to the noisy template model improved our understanding of the *simultaneous* contribution of the bias, sensitivity and similarity between letters in the letter identification task.

## Introduction

By their nature, letters have a complex structure and many studies showed that the legibility is different for different letters ( Sloan, 1959; Ferris et al.,1982; Grimm et al., 1994; Alexander et al., 1997; Reich & Bedell, 2000; Shah et al., 2012; Hamm et al., 2018; Ludvigh, 1941; Strasburger et al., 2011; Hairol et al., 2015; Anderson and Thibos, 2004; Shah et al., 2011). The factors affecting legibility of a given letter are perceivability, similarity with other letters, and the bias towards or against it (Mueller & Weidemann, 2012). The perceivability is a measure of how legible the letter is depending solely on the characteristics of the letters, such as the letter size, contrast, or shape. To avoid ambiguity in using the term perceivability, we will use the term sensitivity instead, which is more relevant in the light of signal detection theory employed here. Response bias is defined as the tendency to favor one response over the other alternatives (Macmillan & Creelman, 1990), and similarity is defined as the confusion in letter recognition which arises among certain letters (e.g., C and O, or N and H). The letter identification could hence be affected by change in sensory input, e.g., size and contrast (sensitivity), the bias towards certain letters in case of uncertainty (response biases), and the confusion between similar letters. Note that from these definitions it is well understood that response biases, unlike sensitivity and letter similarities, are independent of the stimulus.

Luce’s model (1963) has been frequently employed to disentangle the role of similarity and bias on letter identification (Townsend, 1971 a & b; Gilmore et al., 1979; Mueller & Weidemann, 2012). The procedure starts by collecting response data in a letter identification task at the threshold level. The data is collated as a confusion matrix of the presented *vs* responded letters where each cell represents the number of times a given letter was chosen in response to the letter presented. The model is used to decompose the confusion matrix into a bias vector and letter similarity matrix. The bias vector contains the bias parameter for each letter used as a response alternative in the experiment where the sum of all bias parameters values results in the guessing rate of the task. In other words, the response bias of a given letter is the difference between letter usage and guessing rate (Luce, 1963; Mueller & Weidemann, 2012). The similarity matrix shows the similarity parameters for each pair assuming that the similarity is symmetrical (i.e., if H is similar to K, then K is similar to H by the same amount). However, Luce’s choice model does not account for the sensitivity as it is assumed that the sensitivity of a given letter is the average of Luce’s similarity parameters of each letter with the other letters (Mueller & Weidemann, 2012).

Pelli et al. (2006) investigated letter identification efficiency for a wide range of observers’ ages and under various conditions. They concluded that consistent letter identification efficiency across a wide range of conditions suggests that the letter identification task is a fundamental visual process (i.e., not a language skill). To identify a letter, the process starts with identifying the *features* of the letters as opposed to whole-letter templates (Pelli et al., 2006). The letter features are detected by a set of feature detectors (i.e., channels) in the visual cortex (Graham, 1989). These features are considered as crucial factors that distinguish letters from each other and therefore are the key factors in letter identification (Townsend, 1971 a & b; Gilmore et al., 1979; Fiset et al., 2008). Based on the same assumption, the similarity between two letters can be estimated by determining the common features of the two letters. Here, we assume that the similarity between two letters is determined by the correlation features between the letters (Pelli et al., 2006; Fulep et al., 2017). Therefore, letters that share common features (e.g., C and O) would show a higher correlation (i.e., similar letters) compared to the letters with less common features (e.g., V and D; dissimilar letters).

It is important to incorporate the similarity between letters in letter identification models. The challenge is how to incorporate the three factors (bias, sensitivity and similarity) simultaneously in a single model. Mueller and Weidemann (2012) introduced a model to estimate the role of similarity in addition to sensitivity (referred to as perceivability) and response bias simultaneously. They found that all three factors were important in the letter identification task and a model with all three parameters performed better than the two-factor models that only use bias/sensitivity or bias/similarity (Mueller & Weidemann, 2012). However, their model was devised for a two-alternative forced choice (2 AFC) paradigm for letter identification at threshold level. The 2 AFC differs from the letter naming procedure in that it tends to reduce and, hence potentially underestimate bias (Macmillan & Creelman, 2005). Furthermore, the 2 AFC paradigm only measures the similarity between the two presented alternatives whereas in the letter naming procedure (e.g., single-interval 10 AFC, employed here) similarity can be estimated between the presented and all the other alternatives (e.g., 10 letters). Additionally, letter naming procedures are more intuitive and frequently encountered in daily life.

Previously, we investigated the role of response bias and sensitivity in letter identification tasks by devising a model (Noisy Template Model, NTM) based on the framework of signal detection theory in a single-interval 10 AFC letter identification task (Georgeson et al., 2023). The main results suggest that both the response bias towards or against letters and sensitivity differences between letters influence individual letter identification. Additionally, response bias was the key factor in determining how often the letters were responded in the experiment (either correct or incorrect). However, Georgeson et al. (2023) did not estimate the effect of similarity between letters on letter identification, and it was assumed that the sensitivity is a measure of a letter’s similarity with other letters. The current study is a continuation of this work where we introduce a novel and significant extension to the model which allows us to reveal the joint effect of bias, sensitivity and similarity in a single-interval 10 AFC letter identification task.

## Methods

### Participants

Data were collected from 12 observers (six females, mean age 34.25 ± 6.43 (SD), age range: 27 −45 years) with healthy eyes and normal or corrected to normal visual acuity. Written informed consent was obtained from all observers. The study was approved by the University of Plymouth Ethics Committee. The study was conducted in accordance with the Declaration of Helsinki.

### Stimuli

The stimuli used in this study are similar to the ones used in our previous work (Barhoom et al., 2021; Georgeson et al., 2023). We used black letters (luminance = 2.2 cd/m^2^) on a white background (luminance = 215 cd/m^2^), resulting in 99% Weber contrast. A total of 10 standard Sloan letters (C, D, H, K, N, O, R, S, V, Z) were employed. The structure of the letters follows the Sloan letter design so that their height is equal to their width and five times the stroke width.

### Apparatus

The stimuli were presented on a laptop monitor (MSI, GS76 Stealth) with a resolution of 2560 × 1440 and a refresh rate of 240 Hz. Observers viewed the targets at a viewing distance of 70 cm, while sitting on chair without using a chin or forehead rest. The examiner monitored viewing distance by regular checks. At this viewing distance one pixel subtended a visual angle of 0.73 min of arc (′). Experiments were carried out under a room illumination of ∼160 lux. The observer responded by calling out the responses. The responses were entered by the experimenter via a standard computer keyboard to minimize errors caused by mistyping and to improve fixation compliance. The fixation compliance was observed by the experimenter who sat opposite of observer.

### Software

*MATLAB* 2024a (MathWorks, Natick, Massachusetts, USA) was used to implement model fitting and statistical analysis. Routines from the Psychtoolbox-3 were used to generate and present the stimuli (Brainard,1997; Pelli, 1997; Kleiner et al., 2007).

### Procedure

#### Data collection

The experiment was conducted monocularly. The eye was chosen at random. The fellow eye was occluded using an eye patch. The method of constant stimuli was used to collect the data. The letter stimuli were presented for 250 ms and accompanied by an auditory signal. During the experiment, the observers were asked to fixate on a fixation cross (dimensions: length and width = 11.68’, stroke width = 1.46’) presented at the center of the screen. The letters were presented at one paracentral location at an eccentricity of 7 deg from fixation (temporal visual field). Since the variable is the letter size, the peripheral location was chosen (instead of central location) to avoid presenting letter that are too small, potentially exceeding the resolution capacity of the monitor used in the experiment. Multiple pilot experiments were conducted to establish appropriate letter sizes to cover the whole range of responses (ranging from guessing (10%) to certain decisions (100%)). The experimenter entered the responses via a separate computer keyboard connected to the laptop. Five different letter sizes (always defined by their stroke width in minutes of arc, and spaced logarithmically) were tested; 1.50’, 2.20’, 3.24’, 4.76’, and 7’. At each size, each letter was presented 10 times. The Sloan letters with the different identities and sizes were presented randomly in an interleaved design. Each observer completed 500 trials (10 Sloan letters × five letter sizes × 10 presentations per letter). The data were collected in the form of the frequency of responses of each letter to the presented letters (either correct or incorrect). Consequently, for each observer, a confusion matrix of the presented *vs* responded letters was obtained. Only choices of the 10 Sloan letters were accepted. To familiarize the observers with the Sloan letters, the experimenter demonstrated the Sloan letters at the beginning of the session. In the rare cases where observers responded with other letters, the experimenter reminded the observer by presenting the randomized Sloan letters set on the screen. The experimenter prompted a second response from the presented Sloan letter set. After the observer made the choice, the experimenter ensured that the observer had re-fixated before resuming the experiment. Observers showed excellent compliance in responding from the Sloan letter set (on average less than six errors per observer in 500 trials).

### The noisy template model (NTM)

#### Outline of the model

The model presented here is an extended version of our previous NTM (Georgeson et al., 2023) (Figure 2). Previously, we made two simplifying assumptions about the templates. Firstly, there are as many letter templates as there are test letters in the experiment (i.e., 10 templates). Secondly, the templates are assumed to give a response only to their own preferred letter, with no response to other letters (Figure 3 A); that is, the templates are orthogonal. However, in the current version of the model, the second assumption of the NTM (i.e., the orthogonality assumption) has been changed as follows to account for similarity between the letters: the templates respond not only to their preferred letter but also to other, similar letters. That is, the templates are correlated (i.e., non-orthogonal). When a given letter is presented, the output of each letter’s template (i.e., template responses in Figure 2) is subject to bias, sensitivity, and/or similarity to the presented letter. Furthermore, the net output of each template is perturbed by additive Gaussian noise. The bias affects the mean output level of the template by a constant amount whether the template’s preferred letter is presented or not (Figure 3 B). The sensitivity affects the mean output level of the template when the preferred letter is presented (Figure 3 C). The similarity influences the mean output of the non-preferred letter templates according to their correlation to the preferred letter template (i.e., similar letters) (Figure 3 D).

**Figure 1.**
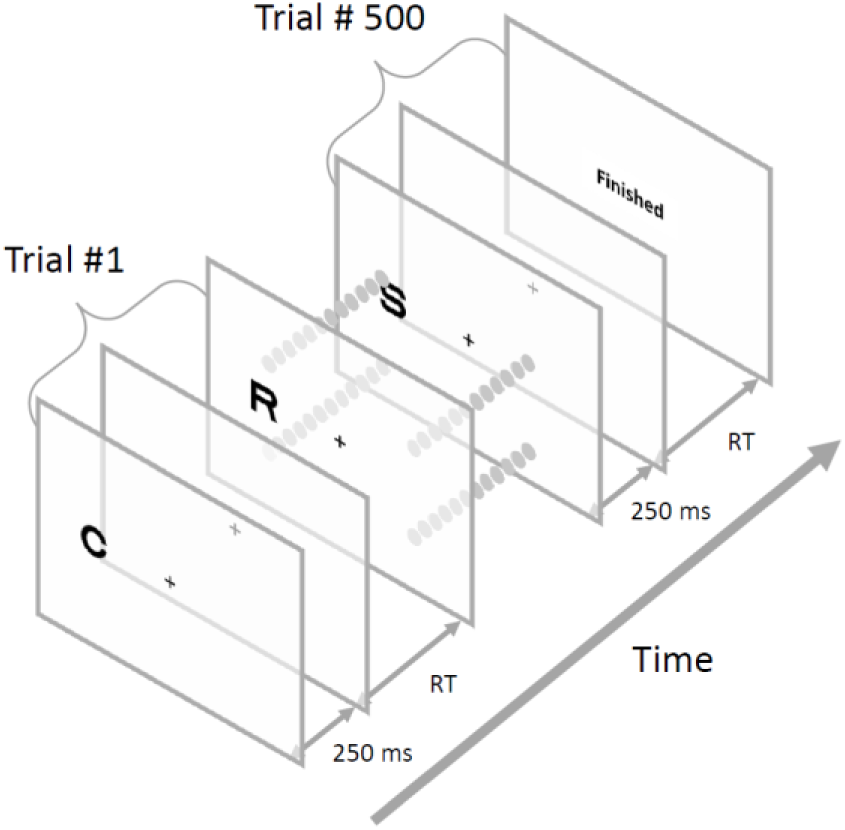
shows the sequence of the events in the trial in the experiment (RT: Response time).

**Figure 2.**
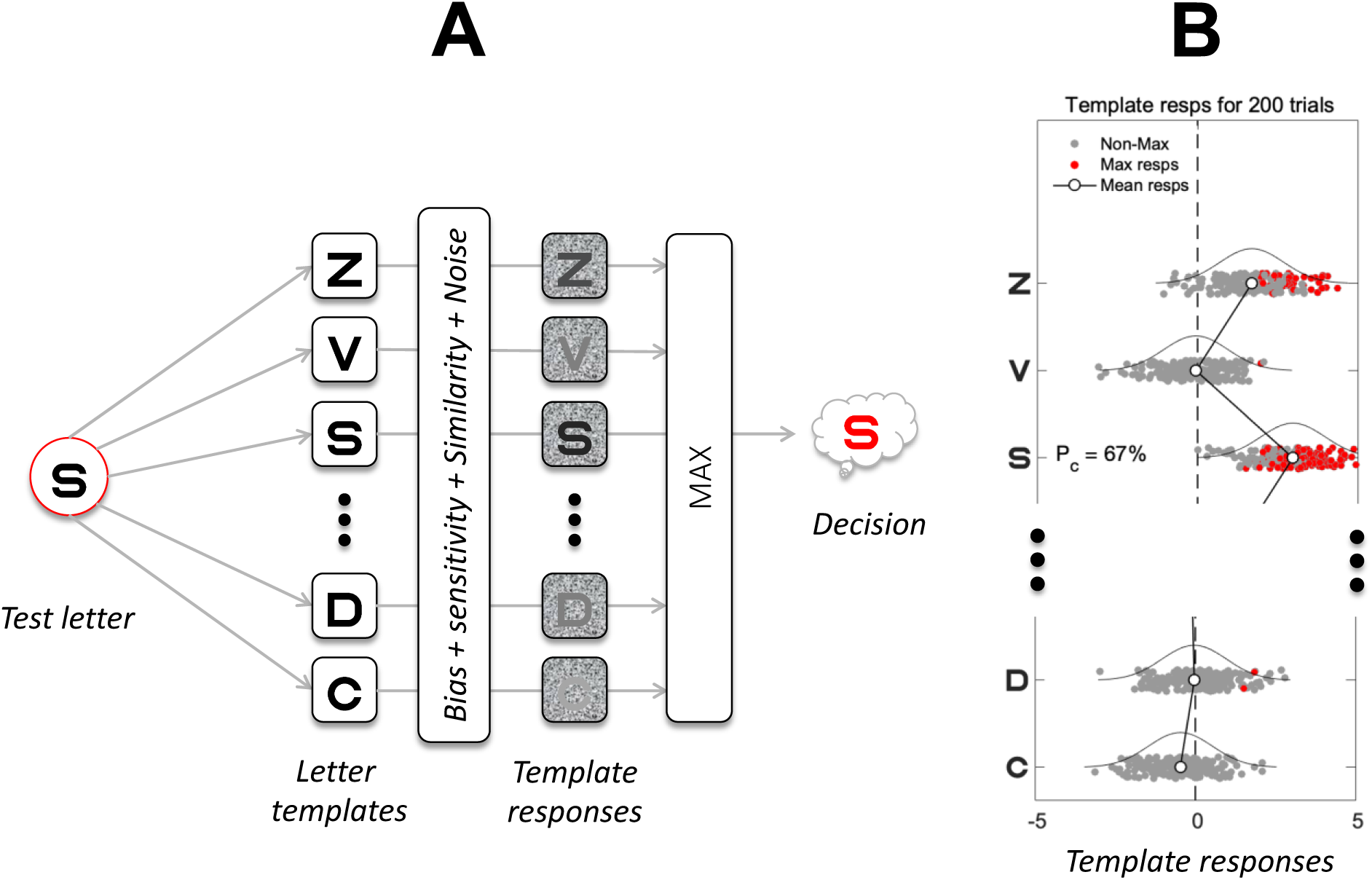
A) a schematic representation of the NTM and (B) template responses. After adding the bias, sensitivity, similarity, and noise on the corresponding templates, the most active (i.e., MAX response) template on a given trial determines the letter choice. For illustration purposes, the red points represent those trials on which a given template gave the maximum response, whereas the grey points are activations that were lower, hence not chosen. Because of noise, the most active template over trials (red dots in B) may be the correct (e.g., S) or the incorrect letter (e.g., Z). The joint effect of the three factors, bias, sensitivity and similarity determines the mean response of each template. For example, positive bias (top three rows) gives a higher mean response compared to negative bias (bottom two rows), hence, increasing the chance of incorrect responses (e.g., red dots for Z) and correct responses (i.e., S). Note that because of the dissimilarity between letter V and S, the mean response of letter V was low (although it has positive bias), therefore, it is unlikely for the template V to be chosen (as an incorrect response in this case). Negative biases (bottom two rows) decrease the chance of these letters being called, correctly or not. However, the similarity between the letters S and D increases the likelihood of letter D (although it has a negative bias) being chosen (as an incorrect response in this case).

**Figure 3.**
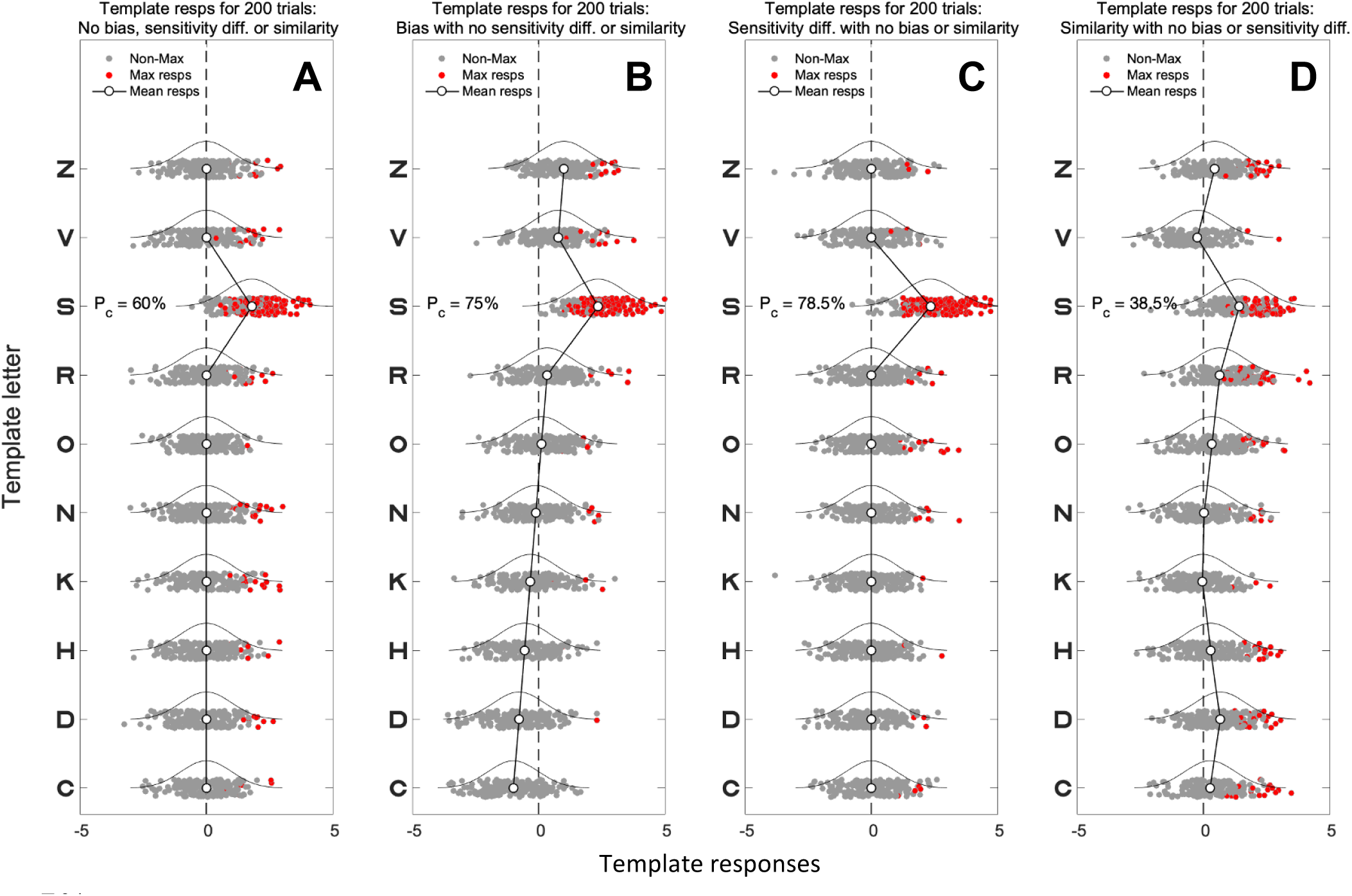
Demonstration of NTM in case of bias, sensitivity, similarity or none. This figure illustrates (A) the NTM without bias, sensitivity differences, or similarity between letters. (B) with different biases on each template. (C) with different sensitivity on each template. (D) with a similarity between templates. The bias and sensitivity in panels B & C are ordered from negative to positive with a mean of zero. We call these the bias gradient B’ and the sensitivity gradient S’ (see text). The letter identity for each template here is arbitrary, for illustration only. Presenting a test letter (e.g., S) increases the mean response for the S-template but leaves others unchanged (A). The letter decision on a given trial is made by choosing the template with the largest response on that trial (the MAX operator, Figure 3 A). Choices vary from trial to trial because of noise in each template channel (indicated by the Gaussian distribution). Notice how the positive bias (e.g., Z), and presentation of the preferred letter (e.g., S) increase the likelihood of choosing certain letters, sometimes correctly, sometimes not (B). In case of sensitivity differences (C), larger sensitivity increases the chance of the letter template to be chosen when its preferred letter is presented (i.e., increase the likelihood of correct responses as shown by the increase of the percent correct response (P_C_) from 60% to 78.5%). Panel D shows the case with no bias or sensitivity differences but with the similarity between the presented letter S and the other templates. Note that the similarity between the presented letter S and the other templates (e.g., D and Z) decreases the likelihood of choosing the letter S (i.e., P_C_ decreased from 60% to 38.5%) and increases the likelihood of choosing the similar letters such as D and Z (i.e., incorrect responses). On the other hand, the dissimilarity between the letters S and V decreases the likelihood of choosing the letter V (i.e., incorrect responses).

### Model structure and equations

For bias and sensitivity modelling, the structure and equations of the original model remained the same as in the original NTM (Georgeson et al., 2023). The current experiment showed a consistent letter usage pattern similar to our previous work (Figure 4). If differences in letter usages are caused by biases, then the most used letter would be associated with the highest positive bias and vice versa. Suppose that the templates can be rank-ordered from least-(most-negative) bias to most-positive bias, indexed by *i* = 1 to *m* (where *m* = 10). Furthermore, bias values *B_i_* (assigned to templates *i* to *m*) are assumed to be a linear function of *i,* ranging from *B_1_* = *-B’* to *B_m_* = *B’*. In this case, the number of free parameters per observer becomes one (i.e., bias parameter (*B’*)) instead of nine.

**Figure 4.**
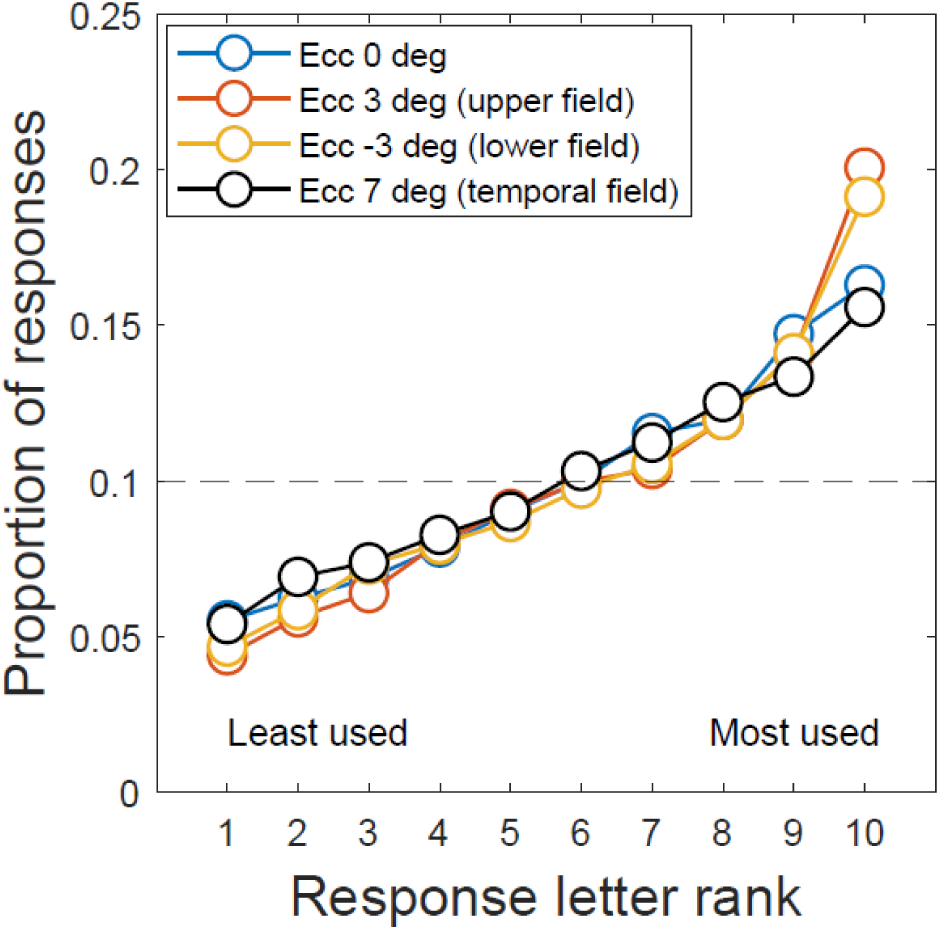
letter usage in the current (Ecc 7 deg) and our previous experiment (Ecc 0, 3 and −3 deg) (Georgeson et al., 2023). Letter usage is expressed as proportion of responses, on which each of the 10 letters was reported (correctly or not), averaged over the letter sizes and observers. The proportion of responses were rank-ordered separately for each observer, then averaged over observers. Note that the gradient of the letter usage of the current experiment is consistent with that of the previous work.

Therefore, the bias *B_i_* for the *i^th^* template is

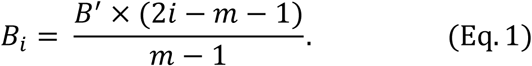

where *B’* is a free parameter. Subsequently, the bias values *B_i_* are assigned to the corresponding templates according to their letter usages. Therefore, the highest bias value is assigned to the template of the most used letter and vice versa. The same procedure is followed to assign the rest of the bias values to the corresponding templates. Note that when *B’* = 0, the system is unbiased.

Similarly, we assumed that sensitivity differs (a linear variation) between templates around a baseline value (i.e., overall sensitivity *S_0_* which is a free parameter) ranging from *S_0_* × (1 *-S’*) to *S_0_* × (1 *+ S’*) where *S’* is the sensitivity gradient. Therefore, the sensitivity of the *i^th^* template to the *j^th^* test letter is

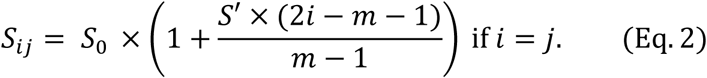

In the model, we refer to *B’* and *S’* as the bias gradient and sensitivity gradient respectively. When fitted to data, *B’* and/or *S’* were free parameters for individual observers, and if not fitted then *B’*=0 and/or *S’*=0. *S_0_* was also a free parameter for each observer, controlling the overall level of performance.

For the similarity simulations and modelling, we assumed that the similarity between Sloan letters arises solely from the features of the letters. Hence, the similarity between pairs of Sloan letters can be quantified by the Optotype Correlation (*OC*) matrix shown in Figure 5 (Fülep et al., 2017). The calculation is based on Pearson’s normalized cross-correlation of the original letters (Neto et al., 2013). To simplify the model, we proposed that similarity is a symmetrical relation (e.g., if H is similar to K, then K is similar to H by the same amount). We also propose that the pattern in the *OC* matrix does not change for different letter sizes.

**Figure 5.**
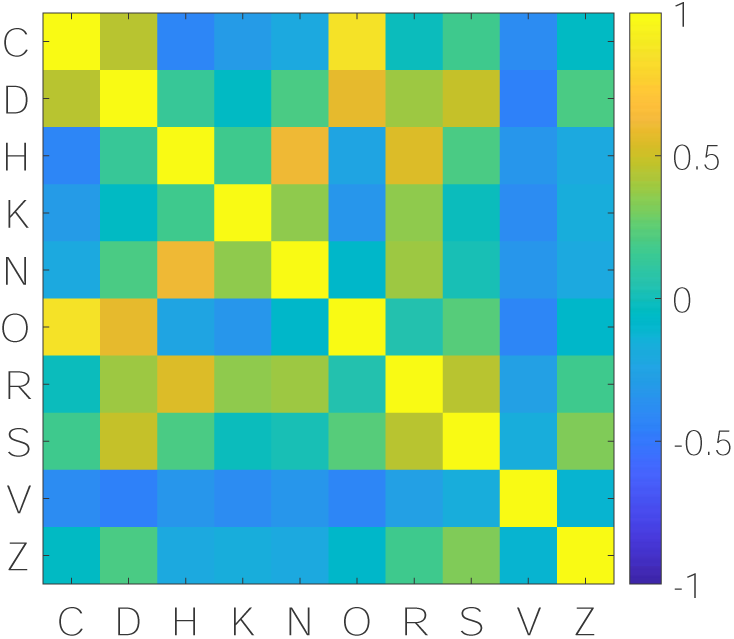
The optotype correlation matrix (*OC*) for Sloan letters. A correlation value of one indicates identical letters, zero means no correlation, and a negative correlation indicates dissimilar letters. A higher positive value means stronger similarity between the corresponding letters.

Another key assumption is that, while keeping the similarity pattern unchanged, the overall strength of the similarity between letters can vary from observer to observer. This is controlled by multiplying the off-diagonal correlation values by a factor that we call the Confusion Strength (*Cs*). Using the *OC* values and the *Cs* factor to quantify the similarity between Sloan letters for a given observer, the sensitivity of the *i^th^* template to the *j^th^*test letter is

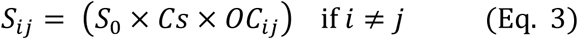

where *Cs* is a free parameter. Thus, the sensitivity of the template *i* when 𝑖 ≠ 𝑗 is controlled by its correlation with the letter j (i.e., *OC_ij_*, see Figure 5) and by *Cs* for a given observer. For example, consider the test letter C and suppose *S_0_* = 1 and *Cs* = 1. In this case, the sensitivity of the letter C template to the test letter C is 1, while sensitivity of the letter O template is 0.866, but that of the (rather dissimilar) letter H template is −0.408. Similar considerations apply to the rest of the templates.

Letter identification improves with increasing letter size. We capture this essential behaviour by assuming that the template’s mean response (*μ_ij_*) increases as a function of sensitivity *S_ij_*, bias *B_i_*, and test letter size *t* (expressed as letter stroke width, in min arc)

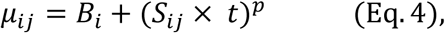

where *p* is an exponent of a power function relation between mean response *μ* and letter size *t*. The exponent *p* controls the slope of the underlying psychometric function (proportion of correct responses *vs* test letter size), unlike our previous model, where *p* was constant. In the current model we set the exponent *p* as a free parameter for each observer. We adopted the standard SDT assumption that the *i^th^*template output is perturbed by additive zero-mean Gaussian noise with variance 𝜎_i_^2^, and that 𝜎_𝑖_ = 1 for all *i*. The decision rule for letter identification was the MAX operator, i.e. the template that has the highest activity is chosen as the response letter (Figure 2 A).

### Letter identification

In the modelling of bias and sensitivity, a more positive 𝜇_𝑖𝑗_ increases the likelihood of the corresponding letter to generate the MAX response (Figure 3 B & C). On the other hand, in the similarity modelling, when a test letter is presented, the mean response becomes more positive for the test letter template but also for similar letters. They will be more likely to generate the MAX response across templates, and tend to be *confused* with the test letter. For letters dissimilar to the test letter, the templates’ mean responses shift in the opposite direction, making them less likely to be confused with the test letter. For example, similar letters like S and D are likely to be confused with each other, whereas dissimilar letters S and V are unlikely to be confused (Figure 3 D). Note that in Eq. 3, if *Cs* is zero, 𝑆_𝑖𝑗_ becomes zero when 𝑖 ≠ 𝑗. This means that our original NTM presented in Georgeson et al. (2023) is a special case of the present model, with zero *Cs*, hence orthogonal templates.

The *MATLAB* function used to compute (for all *i, j*) the probability *P_ij_* that the *i^th^* template delivers a response to the *j^th^* test letter that is larger than that of all the other templates were taken from Zhou et al. (2014). For a given set of parameters (*S_0_*, *p*, *B’*, *S’*, *Cs*), we used this function to compute the matrix of presented *vs* response probabilities from which we calculated the proportions of responses of each letter (total, correct or incorrect) at different letter sizes. i.e., we obtained one 10 × 10 confusion matrix (test letters *vs* response letters) for each tested letter size.

### Illustration of the effect of bias, sensitivity, and similarity on letter identification task

To illustrate the expected effect of bias, sensitivity, and similarity between letters on the response pattern for individual letters, we ran simulations of letter identification tasks. We set 𝑆_0_ = 0.5 and *p* = 1.5 to obtain a reasonable performance for the same five letter sizes tested in the current experiment. The simulations were performed to collect data of the task in case of bias (*B’* = 1, *S’* = 0 and *Cs* = 0), sensitivity (*B’* = 0, *S’* = 1 and *Cs* = 0), similarity between letters (*B’* = 0, *S’* = 0 and *Cs* = 1) or none (*B’* = 0, *S’* = 0 and *Cs* = 0) of the factors incorporated. The bias was simulated so that the *B’* = 1 (chosen for illustration). The calculated biases for individual letters (using Eq. 1) ranged from −1 to 1 and assigned to the letter templates V, N, C, O, H, K, S, D, R and Z (arbitrarily ordered for demonstration), respectively. In this case, the bias of the letter V is −1 (i.e., the largest bias against the letter) and that of the letter Z is 1 (i.e., the largest bias towards the letter). The sensitivity of each letter when its preferred letter is presented (i.e., 𝑖 = 𝑗) was simulated so that *S’* = 1. The calculated sensitivity for individual letters ranged from 0 to 1 (calculated using Eq. 2) and assigned to the same order of letters V, N, C, O, H, K, S, D, R and Z, respectively. In this case, the letter V has the lowest sensitivity whereas the letter Z has the highest sensitivity towards its preferred letter.

The similarity between letters was simulated with *Cs* = 1. The sensitivity of each letter to the non-preferred letter presented (i.e., 𝑖 ≠ 𝑗, hence similarity) is calculated using Eq. 3.

In each scenario, the confusion matrix of the simulated data pooled across the five letter sizes, the proportion of total responses, the proportion of correct responses. and the proportion of incorrect responses were calculated and are depicted in Figure 6. The proportion of total responses for a given letter is calculated as the probability that the letter was reported, whether correct or incorrect, relative to the total number of responses (i.e., 500). The proportion of total responses were then rank-ordered from the lowest (corresponds to the least used letter) to the highest (corresponds to the most used letter) proportion of total responses and are plotted in Figure 6 B. The proportion of correct responses for a given letter is calculated as the probability of calling that letter correctly relative to the total number of correct responses of all letters (Figure 6 C). Likewise, the proportion of incorrect responses for a given letter is calculated as the probability of calling that letter incorrectly relative to the total number of incorrect responses of all letters (Figure 6 D). In the three cases, the chance proportion became 0.1. The order of proportion of correct responses in Figure 6 C and incorrect responses in Figure 6 D follows the same order of the rank-ordered proportion of total response in Figure 6 B.

**Figure 6.**
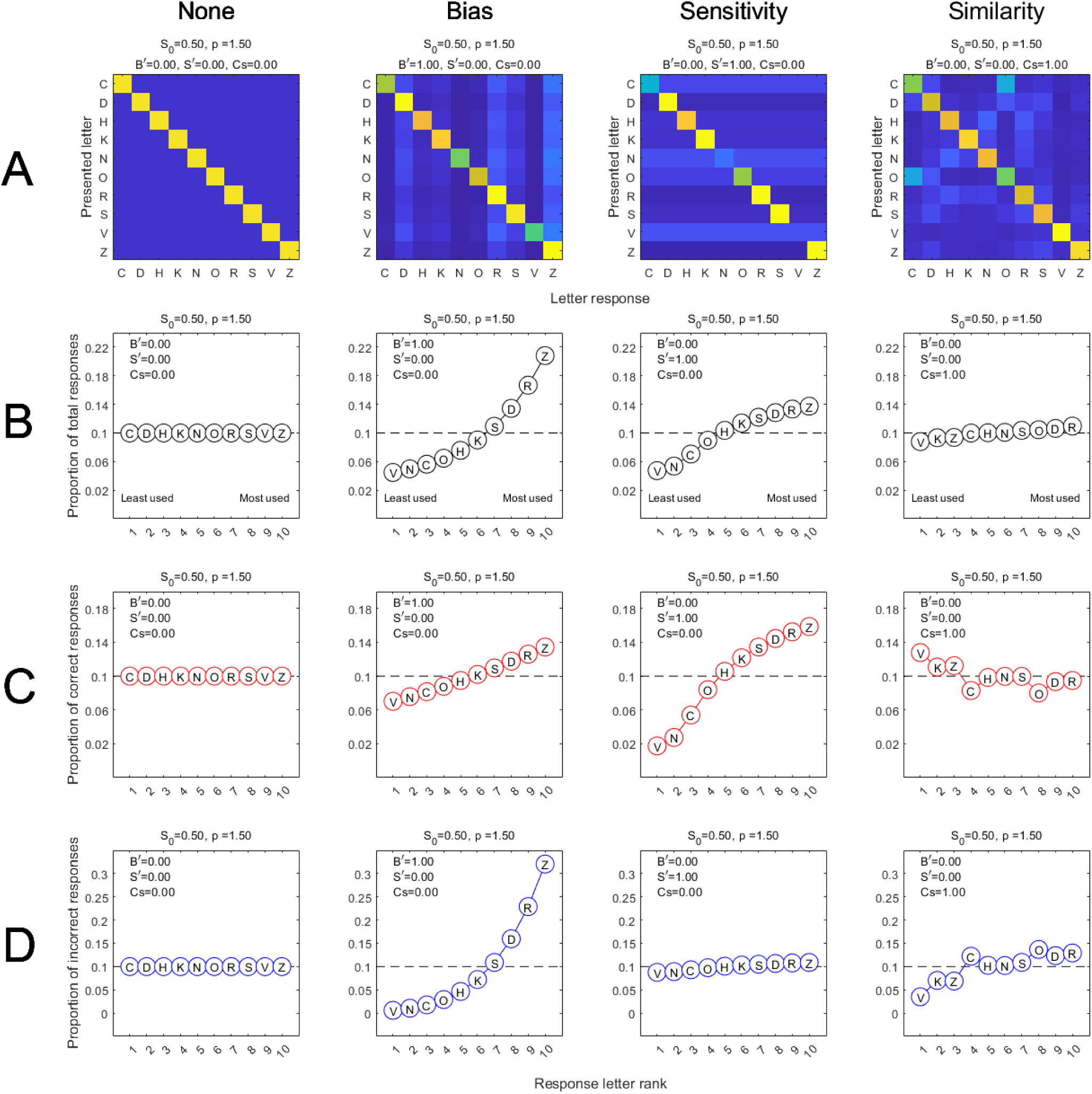
Simulation of the effect of bias, sensitivity, similarity or none on letter identification task. A) the confusion matrix of the simulated data pooled across letter sizes, B) the proportion of total responses, C) the proportion of correct responses and D) the proportion of incorrect responses for each case.

In the case when none of the three factors are incorporated in the simulation (Figure 6, None), due to the absence of bias, sensitivity and similarity between letters, the identical proportion of correct (Figure 6 C, None) and incorrect (Figure 6 D, None) responses across the letters resulted in, as expected, the observed identical proportions of total responses of all letters (Figure 6 B, None). In other words, there is no over-calling or under-calling of any of the letters.

However, in the bias scenario (Figure 6, Bias), the biases added to the letter templates shifted the corresponding proportions of total responses systematically (Figure 6 B, Bias) due to the shifted corresponding proportions of correct and incorrect responses. For example, the simulation shows that the letter Z (assigned the largest bias towards the letter) is over-called (either correctly of not) as shown by the corresponding high proportion correct and incorrect responses. Consequently, the letter Z showed the highest proportion of total responses. The opposite is observed for the letter with the largest negative bias (i.e., V) (Figure 6 C&D, Bias).

In the sensitivity simulation (Figure 6, Sensitivity), the sensitivity differences between letters only affect the proportion of correct responses (Figure 6 C, Sensitivity). In this case, over-calling a given letter (e.g., Z) is due to the increased sensitivity towards that letter when presented (i.e., high proportion correct responses) compared to the other letters. Letter V that is assigned with the lowest sensitivity towards its preferred letter showed the lowest proportion of correct response compared to other letters and therefore the least used.

Figure 6 also shows the performance with only similarity incorporated (Figure 6, Similarity). The simulation shows that the similarity between letters has minimal effect on the proportion of total responses compared to the effect of bias and sensitivity (Figure 6 B, Similarity). This can be explained by the cancellation that occurs between the correct and incorrect responses. For instance, the similarity between two frequently confused letter (e.g., C *vs* O) reduces the proportion of correct responses and increases the proportion of incorrect responses of both letters (Figure 6 C&D, Similarity). Additionally, the more dissimilar a letter is from the rest (e.g., V), the higher the proportion of correct responses and the lower the proportion of incorrect responses for that letter (Figure 6 C&D, Similarity).

### Model fitting

For each observer, we obtained a 10 × 10 (presented *vs* responded) confusion matrix for each of the five letter sizes. The model was fitted to the observed number of responses (i.e., 500 responses), using maximum likelihood–adjusting parameter values *S_0_*, *p*, *B’*, *S’*, and *Cs* to maximize the log-likelihood (*LL*) of the parameters given the data. To calculate the *LL*, we used the following equation:

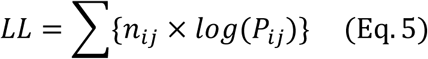

where *n_ij_* is the observed number of responses of the letter *i* when presenting the letter *j*. *P_ij_* is the model’s proportion of responses for the corresponding *ij* pair. The variables *n* and *P* range over 100 letter pair combinations for each letter size, and the summation takes place over those 500 pairs.

We fitted eight versions of the template model, in which the fitted parameters represented *B’*, *S’* & *Cs*, denoted as the model (B1C1S1), or *B’* & *Cs* (B1C1S0), *B’* & *S’* (B1C0 S1), *Cs* & *S’* (B0C1S1), *B’* only (B1C0S0), *Cs* only (B0C1S0), *S’* only (B0C0S1) or none (B0C0S0). The fitting of the B1C1S1 model was performed in two steps. Firstly, we adjusted overall sensitivity *S_0_* and slope (*p*) in a 2D sampled grid search to find the best-fitting value that maximised *LL* assuming that there is no bias (*B’* = 0), no sensitivity differences (*S’* = 0), and no similarity between letters (*Cs* = 0). Secondly, while using the best *S_0_* and *p* we ran a 3D sampled grid search to find the best values of the *B’*, *Cs*, and *S’*. Then we readjusted for *S_0_* and *p* using the best *B’*, *Cs*, and *S’*. We repeated the procedure until there was no change in the estimated parameters.

A similar approach was used to fit the B1C1S0, B1C0S1, and B0C1S1 models. In these cases, a 2D sampled grid search was performed first to fit for *S_0_* and *p*, followed by 2D sampled grid search to fit for either *B’* & *Cs*, *B’* & *S’* or *Cs* & *S’*. We repeated the procedure until there was no change in the estimated parameters. For the remaining models, a 3D sampled grid search was performed to fit the B1C0S0, B0C1S0, and B0C0S1 models to find the best values of the (*S_0_*, *p* & *B’*), (*S_0_*, *p* & *Cs*) or (*S_0_*, *p* & *S’*) respectively. A 2D sampled grid search was performed to fit the B0C0S0 model to find the best value of the *S_0_* and *p*. All the fittings were performed for each observer separately.

In the following section, we present the results of fitting the eight models to the experimental data and the comparison between them via Akaike Information Criterion (AIC, Akaike, 1974) to determine the best fitting model.

## Results

### Initial observations

The results of the experimental data are summarized in Figures 7 and 8. Figure 7 A shows the confusion matrix (presented *vs* responses of letters) collapsed across observers and letter sizes. The proportion of total, correct and incorrect responses in Figure 7 B, C&D respectively, show the characteristic systematic shift in performance. Through simulation, we can demonstrate that the same systematic shift in performance occurs when incorporating bias and sensitivity in the task (see “Illustration of the effect of bias, sensitivity and similarity on letter identification task” in methods). Figure 8 A shows the confusion matrix of each letter size averaged across observers. Note that, the pattern of responses at larger letter sizes (e.g., letter size #3) resembles the pattern of the OC matrix (Figure 5), suggesting that the performance is also influenced by the similarity between letters. Since the data here is averaged across observers, and each observer might show different preference in biases or sensitivities towards letters, the effect would be diminished by averaging. Figure 8 B shows the confusion matrices recalculated as the average across the rank-ordered letter responses according to the rank of the overall letter usage (i.e., the rank of the letters total responses pooled across all letter sizes) of each observer. This reveals that the observers clearly overcall some letters over the others, especially at the two smallest letter sizes. The maximum effect is at the smallest letter sizes where the responses will be determined mainly by bias. Note that at the two smallest letter sizes, where there is high uncertainty in the performance, the responses are determined mainly by the bias which is, unlike similarity and sensitivity, independent of the sensory input (i.e., letter size). As the letter size increases, the effect of similarity (frequency of responses of certain letter pairs) and/or sensitivity (frequency of the correct responses, i.e., diagonal cells) become more prominent. Different model variants were fitted to the experimental data to reveal the effect of the three factors (i.e., bias, sensitivity and/or similarity) in the letter identification task in the current experiment (Figure 9).

**Figure 7.**
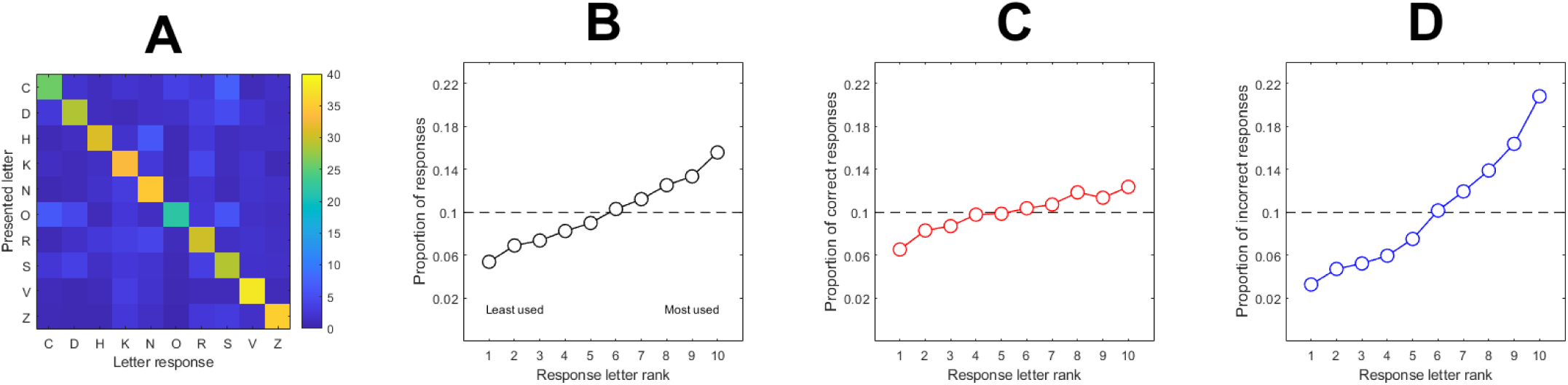
shows the experimental data as A) confusion matrix (presented *vs* responses of letters). Data is collapsed across letter sizes first and then averaged across observers. Here and throughout (unless mentioned otherwise), the colour scale illustrates the frequency of letter responses, where warmer colours show higher frequencies. The diagonal cells represent correct responses, whereas the non-diagonal cells represent incorrect responses. The figure also shows B) the proportion of total responses, C) the proportion of correct responses and D) the proportion of incorrect responses. The letters were rank-ordered according to their proportion of total responses for each observer from the least to the most used letters then averaged. The proportion of correct and incorrect responses follow the same order of the proportion of total responses.

**Figure 8.**
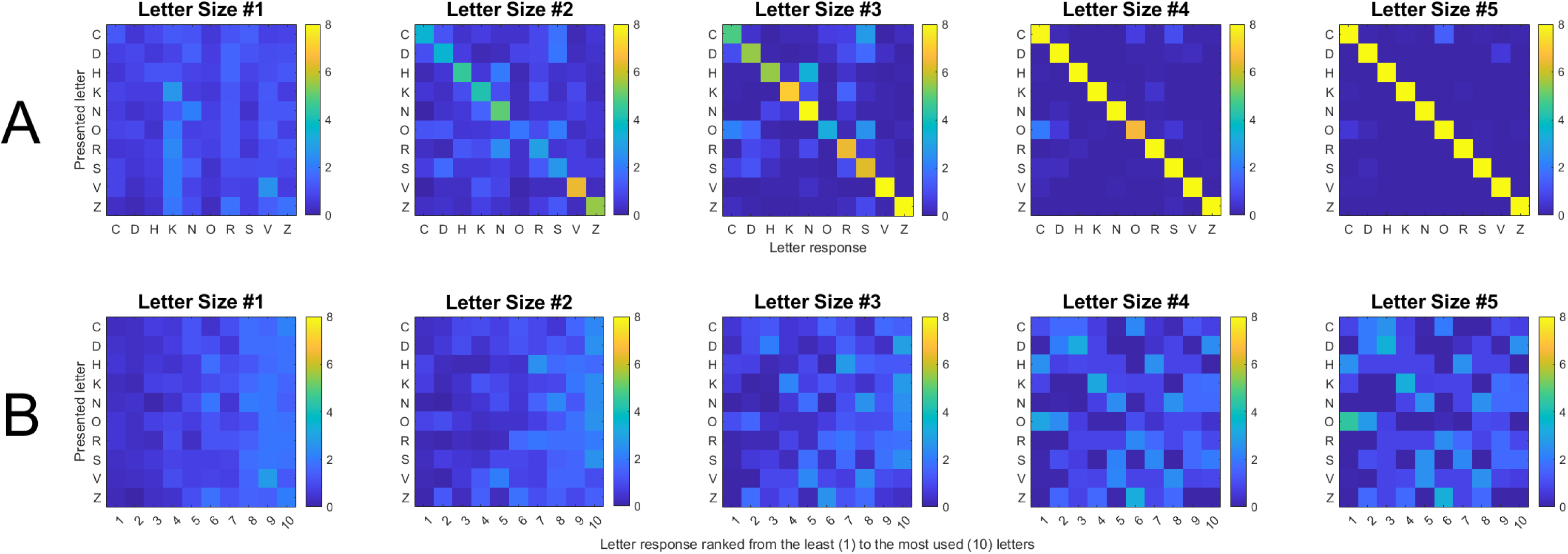
shows A) the experimental data presented as the confusion matrix of each letter size averaged across observers. B) Shows the confusion matrices recalculated as the average across the rank-ordered letter responses (from the least to the most used letters) according to the overall letter usage (i.e., the rank of the letters total responses pooled across all letter sizes) of each observer.

**Figure 9.**
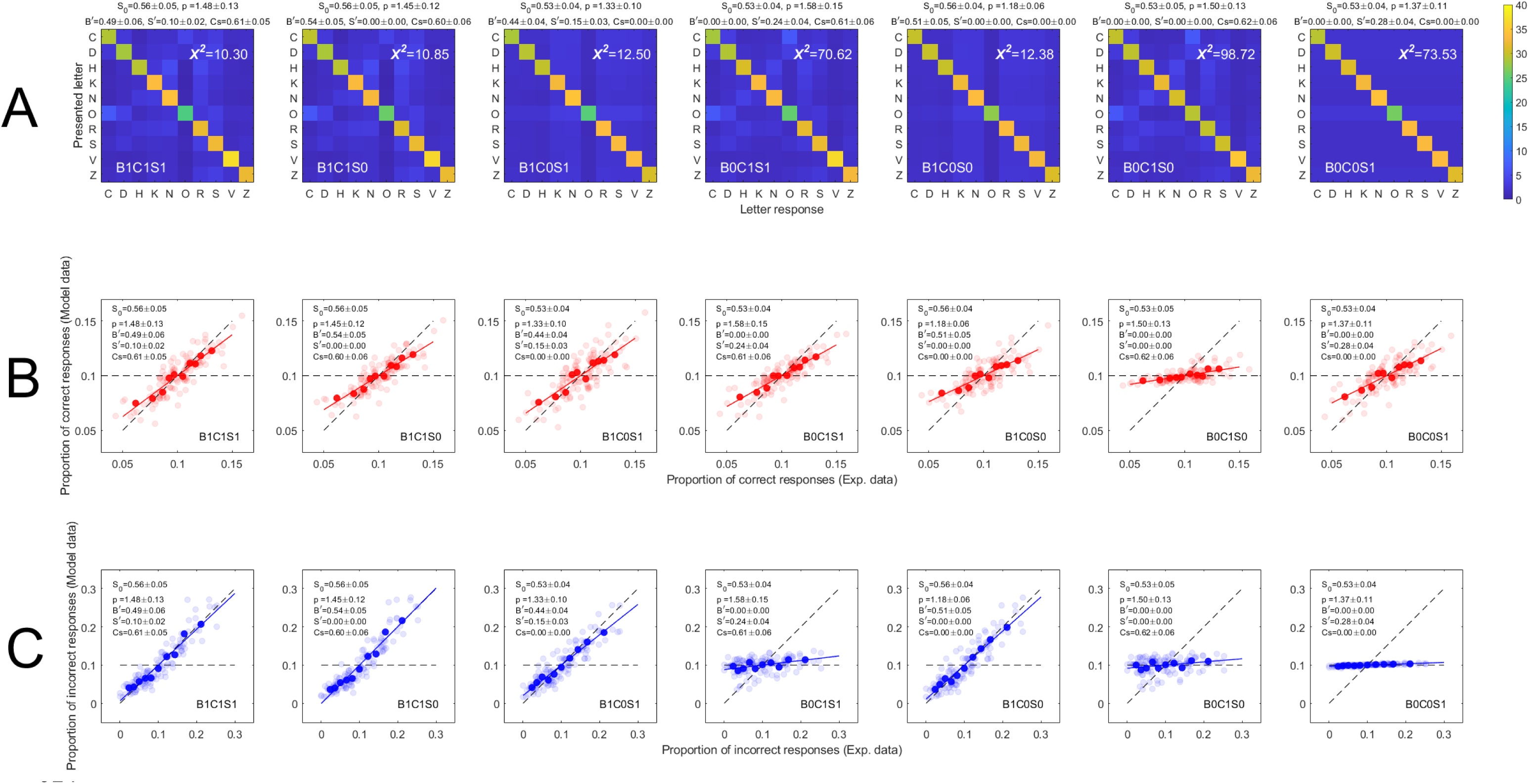
shows the seven variants of the model after excluding the basic model B0C0S0. A) for each model, the confusion matrix is averaged across the five letter sizes for each observer and then averaged across observers (after rank-ordering the data, see text). 𝜒^2^ value in each matrix shows the difference between a given model and the experimental data. The smaller the value, the better the model is in accounting for the experimental data. B) shows the agreement between the model and experimental data in the proportion of correct responses. The faint red points are the observers’ data for individual letters. For each observer, the letters’ proportions of correct responses in experimental data were rank ordered first (from low to high proportions). Then the ranked proportions of the experimental data were averaged across observers. The model data average was calculated across observes’ model data using the same ranking (i.e., based on ranking the proportion of correct responses of the experimental data). The solid red data points (and the fitted red line) show the agreement between the two averages. The same procedure was applied to calculate and depict (using blue colour) the averages of the proportion of incorrect responses (C). Here and throughout (unless mentioned otherwise), the diagonal dashed line represents the perfect agreement whereas the horizontal dashed line represents the complete absence of agreement (i.e., chance performance).

Figure 9 A shows the resulting confusion matrices of the fitted models averaged across the five letter sizes and then averaged across observers, where each column shows the fit of one model (the basic model B0C0S0 is not included). To facilitate the identification of the model that accounts best for the experimental data (Figure 7 A), the difference between each rank-ordered model and rank-ordered experimental data was calculated using the 𝜒^2^value. To minimize the effect of averaging, for each observer, model and experimental data were rank-ordered first according to the total letter usage of experimental data, then averaged. For each model and each location (Figure 9 A), 𝜒^2^ value was calculated as:

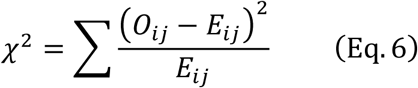

where *O_ij_* is the frequency of responses of the cell *ij* in the confusion matrix of experimental data (i.e., observed) and *E_ij_* is the frequency of responses of the corresponding cell (i.e., *ij*) in the confusion matrix of the model data (i.e., expected).

Inspection of Figure 9 A suggests that the fit of the bias, sensitivity, and similarity model (i.e., B1C1S1) and the fit of the bias and similarity model (i.e., B1C1S0) remarkably resemble the pattern of experimental data (Figure 7 A). Additionally, model B1C1S1 and B1C1S0 show the smallest 𝜒^2^ values of 10.30 and 10.85, respectively. Model B1C1S1 and B1C1S0 also show a good account for the proportions of both correct and incorrect responses at individual and group levels (Figures 9 B & C). Figure 9 also suggests that bias is the most influential factor in model B1C1S1 and B1C1S0 since it has the smallest 𝜒^2^ value (𝜒^2^ = 12.38) when fitted individually (i.e., B1C0S0 ) compared to the sensitivity (i.e., B0C0S1) where value is higher (𝜒^2^ = 73.35), and compared to similarity (i.e., B0C1S0) where value is also higher (𝜒^2^ = 98.72) (Figure 9 A). Furthermore, the bias is the key factor that accounts for some of the proportion correct and most (if not all) of the incorrect responses when fitted individually (i.e., B1C0S0) or in conjunction with the other factors (i.e., B1C1S0, B1C0S1, and B1C1S1) (Figure 9 B & C). Since the 𝜒^2^values are calculated for average across the rank-ordered data, the results here might underestimate the contribution of the similarity factor. However, Figure 9 B&C show that the similarity factor (when fitted individually) poorly accounted for either correct or incorrect responses compared to the sensitivity factor when fitted individually (showed some accountability for the correct responses and none for the incorrect responses) or compared to the bias factor when fitted individually (showed some accountability for the correct responses and most of the incorrect responses).

### Formal model comparison

To test these observations, we conducted a formal model comparison using the *AIC* (Akaike, 1974) at the group level (Table 1). *AIC* weights were 1 for the fuller model B1C1S. Therefore, the fuller model B1C1S1 is the best or most-favoured model among the eight candidates (Table 1).

**Table 1.** Comparison of the eight models via AIC analysis, for the group of 12 observers.

| Model | *LL* | *K* | *n* | *AIC* | $\Delta AIC$ | *Ak. Wt* |
| --- | --- | --- | --- | --- | --- | --- |
| B0C0S0 | -21493.33 | 24 | 6000 | 43034.87 | 1761.30 | 0 |
| B0C0S1 | -21282.15 | 36 | 6000 | 42636.75 | 1363.18 | 0 |
| B0C1S0 | -21137.97 | 36 | 6000 | 42348.40 | 1074.83 | 0 |
| B1C0S0 | -20980.90 | 36 | 6000 | 42034.25 | 760.679 | 0 |
| B0C1S1 | -20967.62 | 48 | 6000 | 42032.04 | 758.47 | 0 |
| B1C0S1 | -20933.68 | 48 | 6000 | 41964.16 | 690.59 | 0 |
| B1C1S0 | -20595.40 | 48 | 6000 | 41287.59 | 14.03 | 0 |
| B1C1S1 | -20576.17 | 60 | 6000 | 41273.58 | 0 | 1 |

Here and throughout: *LL*, total log likelihood; *K*, total no. of free parameters across the group; *n*, total no. of data points; *AIC* is the Akaike scores; the $\Delta AIC$ is the relative difference of the scores; *Ak. Wt* refers to the Akaike weights.

This finding is also confirmed by the analysis of deviance (Collett, 2003). The analysis of deviance showed that bias (B0C1S1 *vs* B1C1S1, *Chi* = 782.90, *df* = 12, *p*<0.001), sensitivity (B1C1S0 *vs* B1C1S1, *Chi* = 38.46, *df* = 12, *p*<0.001), and similarity (B1C0S1 *vs* B1C1S1, *Chi* = 715.02, *df* = 12, *p*<0.001) factors have important role in the fitting of the fuller model B1C1S1.

Further model comparisons after excluding the fuller model (B1C1S1) reveal that the bias and similarity model (B1C1S0) is favoured over the other six models (Table 2). Additionally, comparing only the one-factor models reveals that the bias model (B1C0S0) is favoured over the other two models (B0C1S0 & B0C0S1) (Table 3). Comparing the similarity model to the sensitivity model after excluding the bias model shows that the similarity model is favoured over the sensitivity model (Table 4).

**Table 2.** Comparison of the seven models via AIC analysis, for the group of 12 observers.

| Model | $LL$ | $K$ | $n$ | $AIC$ | $\Delta AIC$ | $Ak.Wt$ |
| --- | --- | --- | --- | --- | --- | --- |
| B0C0S0 | -21493.33 | 24 | 6000 | 43034.87 | 1747.28 | 0 |
| B0C0S1 | -21282.15 | 36 | 6000 | 42636.75 | 1349.16 | 0 |
| B0C1S0 | -21137.97 | 36 | 6000 | 42348.40 | 1060.80 | 0 |
| B1C0S0 | -20980.90 | 36 | 6000 | 42034.25 | 746.654 | 0 |
| B0C1S1 | -20967.62 | 48 | 6000 | 42032.04 | 744.45 | 0 |
| B1C0S1 | -20933.68 | 48 | 6000 | 41964.16 | 676.56 | 0 |
| B1C1S0 | -20595.40 | 48 | 6000 | 41287.59 | 0 | 1 |

**Table 3.** Comparison of the four models via AIC analysis, for the group of 12 observers.

| Model | $LL$ | $K$ | $n$ | $AIC$ | $\Delta AIC$ | $Ak.Wt$ |
| --- | --- | --- | --- | --- | --- | --- |
| B0C0S0 | -21493.33 | 24 | 6000 | 43034.87 | 1000.62 | 0 |
| B0C0S1 | -21282.15 | 36 | 6000 | 42636.75 | 602.50 | 0 |
| B0C1S0 | -21137.97 | 36 | 6000 | 42348.40 | 314.15 | 0 |
| B1C0S0 | -20980.90 | 36 | 6000 | 42034.25 | 0 | 1 |

**Table 4.** Comparison of the three models via AIC analysis, for the group of 12 observers.

| Model | $LL$ | $K$ | $n$ | $AIC$ | $\Delta AIC$ | $Ak.Wt$ |
| --- | --- | --- | --- | --- | --- | --- |
| B0C0S0 | -21493.33 | 24 | 6000 | 43034.87 | 686.47 | 0 |
| B0C0S1 | -21282.15 | 36 | 6000 | 42636.75 | 288.35 | 0 |
| B0C1S0 | -21137.97 | 36 | 6000 | 42348.40 | 0 | 1 |

These findings suggest that bias has a higher contribution to the fuller model than the other two factors (sensitivity and similarity) and the joint effect of bias and similarity is the most influential in the fuller model than the joint effect of either bias and sensitivity or similarity and sensitivity. The results also suggest that the sensitivity factor has the least effect on the fuller model.

### The relative role of bias, sensitivity, and similarity in the letter identification task

The “best” model (i.e., B1C1S1) does not necessarily mean that the model is good at explaining different aspects of the letter identification task in the current experiment. We showed that the best model accounts very well for the differences in the proportion of correct and incorrect responses (Figure 9 B & C). But it is crucial to investigate how well the best model captures the bias, sensitivity, and similarity separately in the experimental data, which we will describe in the following section.

Figure 10 shows the agreement of the best model with experimental data in letters correct (red) and incorrect (blue) frequencies in case of the model recomputed with three factors activated, sensitivity and similarity activated (i.e., B off), bias and sensitivity activated (i.e., C off), bias and similarity activated (i.e., S off), only sensitivity activated (i.e., B & C off), only similarity activated (i.e., B & S off) or only bias activated (i.e., C & S off). Results show that the bias is the key factor in explaining the overall response (Figure 10, Letter Size #1-5) (i.e., the differences in correct and incorrect response frequencies) and the response pattern at each letter size in the experimental data either individually (i.e., C & S off) or in conjunction with the other factors (i.e., B1C1S1, C off and S off) (Figure 10). On the other hand, one would assume that the sensitivity differences between letters would account for the differences in correct responses in the experimental data, but that was not the case when we examined the model recomputed for sensitivity individually (i.e., B & C off). The model with only the similarity factor activated (i.e., B & S off) also showed poor accounting for differences in correct and incorrect responses in experimental data. Furthermore, even the model with the joint effect of sensitivity differences and similarity between letters only (i.e., B off) does not account for responses (especially the incorrect responses) in experimental data as the model with the bias individually does (i.e., C & S off).

**Figure 10.**
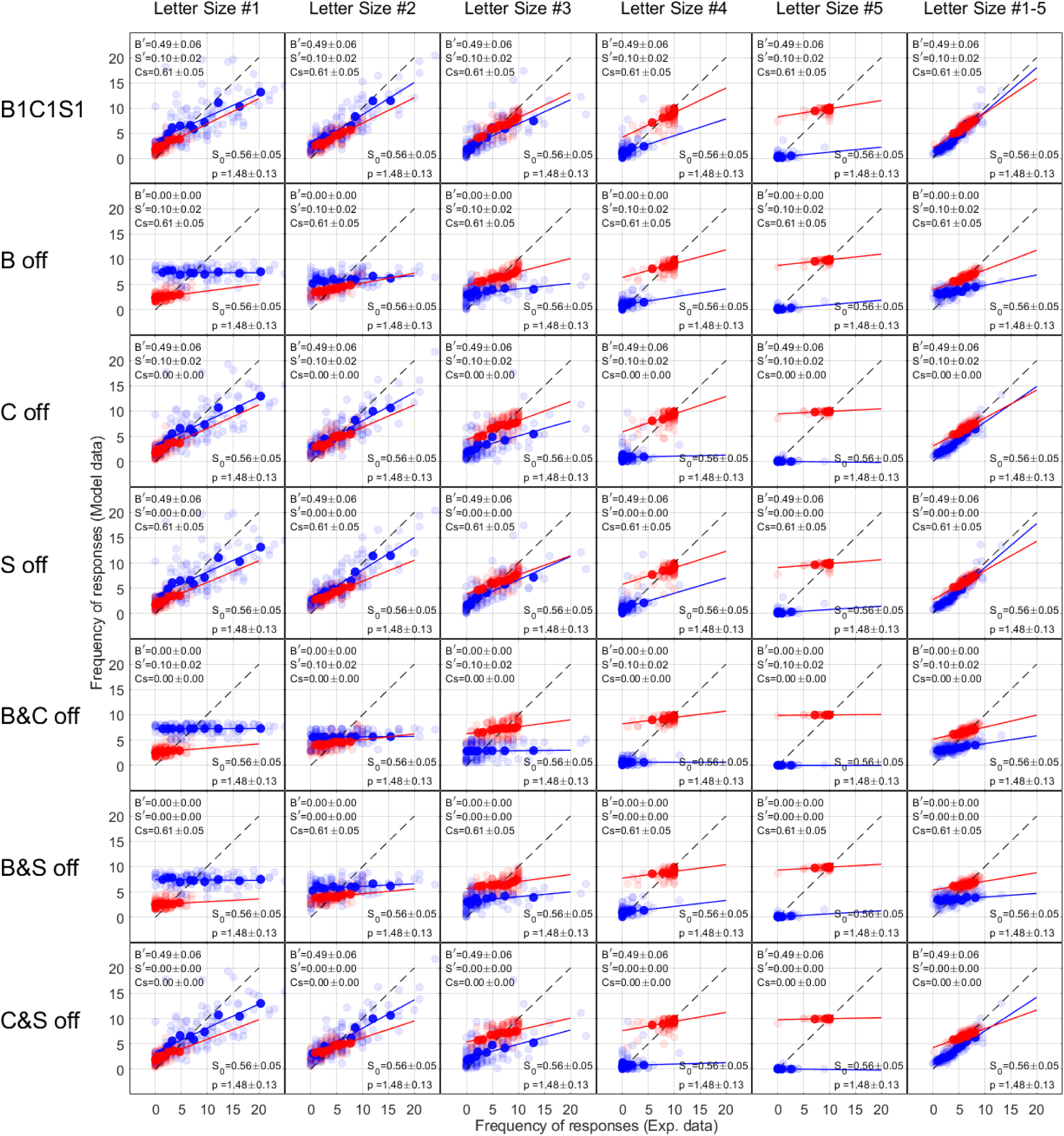
shows the best model (B1C1S1) recomputed with different activated factors (see text). It shows, the agreement between experimental and model data in the differences of letter correct and incorrect response frequencies. Bias is the key factor in defining the overall responses and the differences in letter correct and incorrect responses in experimental data.

Results show that the best model (B1C1S1) accounts for the differences in the proportion of *incorrect* responses (Figure 9 C). However, here we investigated more specifically how the best model accounts for the sources of the differences in incorrect responses which can be either bias, similarity between letters, or both. We employed Luce’s choice model (Luce, 1963) to compute the bias and similarity from the best model B1C1S1 and the experimental data, and then examined the agreement between the results of the model and experimental data (Figure 11). A custom-made *MATLAB* function to compute the response biases and similarities parameters using Luce’s choice model was employed and is freely accessible here: https://github.com/HBarhoom/Codes- . The fitting of Luce’s choice model showed that model B1C1S1 described the bias and similarity in the experimental data very well. Figure 11 A shows that Luce’s bias in the best model is strikingly similar to Luce’s bias in the experimental data. This also indicates that our model estimates the same kind of bias that Luce’s model does, and it supports our strong assumption about the linearity of the bias and its ranking based on the total letter usage. Figure 11 B shows Luce’s similarity parameters of the 45 confusion pairs. There are only 45 confusion pairs because, Luce’s and our model assume that the similarity between letters is symmetrical (i.e., the parameter of similarity between the presented letter C and responded letter O is identical to the similarity parameter between the presented letter O and responded letter C). The results suggest that Luce’s similarity between letters estimated from the best model data is similar to Luce’s similarity between letters estimated from experimental data. It is also found that the agreement is not as strong as in the case of Luce’s bias comparison (Figure 11 B). This is not unexpected since we used the *OC* matrix to capture the similarity between letters that follows the pattern of the *OC* matrix. Hence, Luce’s model will capture a particular pattern of similarity when fitted to the best model data (i.e., the pattern of *OC* matrix) whereas Luce’s model captures different and a variety of similarity patterns for each observer when fitted to experimental data. Nevertheless, the reasonable agreement between Luce’s similarity of the best model and experimental data suggests that the general and common similarity pattern in experimental data is the one that follows the *OC* matrix (Figure 5)

**Figure 11.**
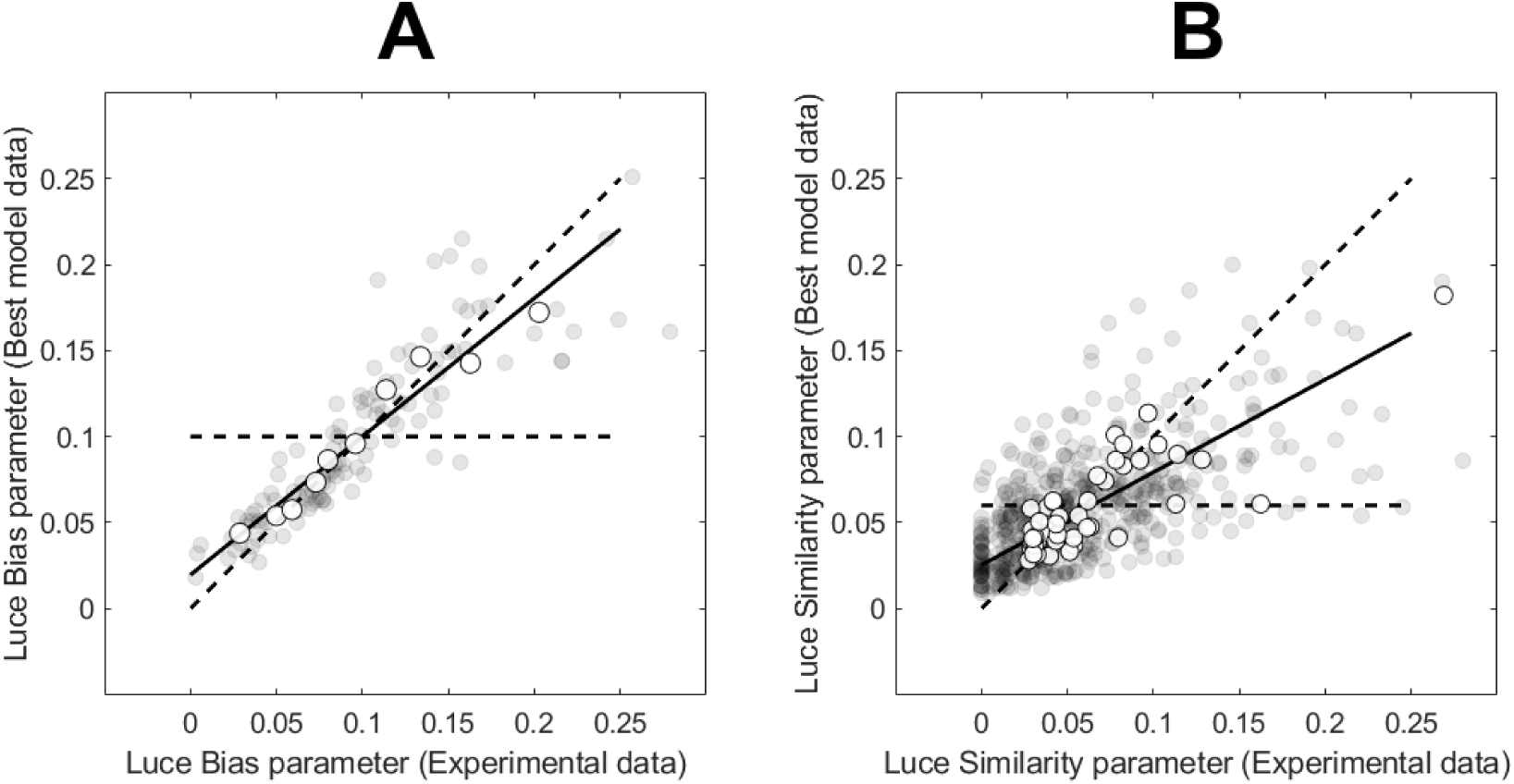
shows A) the agreement between Luce’s bias parameters estimated from the best model and experimental data. B) shows the agreement between Luce’s similarity parameters of the model and experimental data. The faint grey points are the observers’ data. The white points (and the fitted black lines) show the agreement between the averages of the observers’ data.

## Discussion

Georgeson et al. (2023) recently introduced a model to compare the influence of biases and sensitivities of individual letters in letter identification. Here we extended this model by incorporating the similarity between letters in the NTM to reveal the joint role of the three factors bias, sensitivity and similarity (instead of bias *vs* sensitivity only).

The results clearly show that the model with the three factors (i.e., B1C1S1) was the *best* model and gave an excellent account to the total, correct and incorrect responses in the experimental data (Figure 9, Table 1). Results also show that the biases are the major source of the errors (i.e., incorrect response) in the task and contributes to a greater degree than the similarity or the sensitivity do to the differences of the correct responses between letters (Figure 10).

Modelling the bias assumes that the biases to individual letters are rank-ordered based on the overall usage of these individual letters when pooled across the letter sizes. The rank-order is unique for each observer. This assumption has been investigated extensively in our previous work and the experimental data supports such a systematic order (Georgeson et al., 2023).

For the modelling of the similarity, we assumed that the patterns of similarity between letters for a given observer must follow the *OC* matrix pattern. It is very unlikely that the pattern of similarity observed and captured by the model in the experimental data to be result of random error (i.e., noise). We examined this assumption via simulation (see appendix). We showed that the random error alone in the letter identification task is very unlikely to create an *apparent* similarity pattern (*p* < 0.0001).

Luce’s choice model (Luce, 1963) has been shown to perform very well in capturing response bias and similarity in letter identification tasks (Nosofsky, 1991; Smith, 1992; Mueller & Weidemann, 2012; Coates, 2015; Hamm et al., 2018). Here we employed Luce’s choice model to further validate our model. We investigated whether the best-performing model (B1C1S1) is efficient in capturing bias and similarity by calculating and comparing bias and similarity parameters from model B1C1S1 and the experimental data using Luce’s choice model. Results show that model B1C1S1 was remarkably efficient (especially for the bias) in capturing the bias and similarity as shown by the agreement in Luce’s bias and similarity parameters estimated from model B1C1S1 and experimental data (Figure 11).

In the current study, we examined the fitting of the proportions of responses of the model to the corresponding number of responses in the experimental data (either correct or incorrect). This means that we imposed a high restriction on the model fitting so that the correct and incorrect responses in the experimental data must support our strong assumptions regarding letter usage-based rank-order and *OC* pattern of similarity. Interestingly, the model showed an excellent account to the experimental data proving that our assumptions were valid to a large extent.

The current study might underestimate the effect of similarity and its contribution to the *best* models fit. This is because modelling the bias and similarity are essentially different in the sense that modelling the bias uses some information from the experimental data (i.e., the total usage rank-order) whereas the similarity modelling was entirely objective where no information from experimental data were used (i.e., we only used the *OC* matrix as an objective source of information to fit the model to experimental data). One would expect that if some information from the experimental data were used to relax the similarity model by taking into account the individual difference in *OC* patterns (which is highly expected), the fitting of the similarity model would be much better and its contribution to the *best* model B1C1S1 fitting would be higher. In the future it would be desirable to improve the structure of the similarity model to take into account any individual differences in the *OC* pattern.

Nevertheless, incorporating the similarity factor (with the current assumptions) to the NTM improved our understanding of the simultaneous contribution of the bias, sensitivity and similarity between letters in the letter identification task.

## Appendix

1. Does the random error (i.e., noise) in the letter identification task produce an *apparent* similarity pattern?

We examined the hypothesis that the random error alone (i.e., noise only with no similarity, sensitivity or bias) in the letter identification task in the current experiment can produce *apparent* similarity pattern effect in the data. To examine this, we simulated data (trial by trial Monte Carlo simulation using the NTM without similarity, sensitivity or bias) that mimics the letter identification task in the current experiment at the 7deg temporal location (i.e., p = 2.5, overall sensitivity = 1.1, 10 letters, 10 simulations per letter for each letter size of 1.50’, 2.20’, 3.24’, 4.76’, and 7’.). Then, we fit the model B0C1S0 using the same procedure described in the method section (Model fitting) to estimate the confusion strength (*Cs*) in the simulated data. We repeated the procedure 10000 times (i.e., 10000 estimations of *Cs*). Finally, we compared the results to the experimental data. We found that the similarity pattern observed in experimental data was very unlikely to be due to the random error alone (*p* < 0.0001).

## Notes

### Competing Interest Statement

The authors have declared no competing interest.

